# A comparative analysis of label-free liquid chromatography-mass spectrometry liver proteomic profiles highlights metabolic differences between pig breeds

**DOI:** 10.1101/346056

**Authors:** Samuele Bovo, Alessio Di Luca, Giuliano Galimberti, Stefania Dall’Olio, Luca Fontanesi

## Abstract

Liver is a complex organ governing several physiological processes that define biological mechanisms affecting growth, feed efficiency and performance traits in all livestock species, including the pig. Proteomics may contribute to better understand the relationship between liver functions and complex production traits in pigs and to characterize this specie as biomedical model. This study applied, for the first time, a label-free liquid chromatography-mass spectrometry (LC-MS) proteomic approach to compare the liver proteome profiles of two important heavy pig breeds, Italian Duroc (IDU) and Italian Large White (ILW). Liver specimens were collected (after slaughtering) from performance tested pigs of these two breeds, raised in standard conditions. The label-free LC-MS method captured a total of 501 proteins of which 200 were subsequently considered in the between breeds comparison. A statistical pipeline based on the sparse Partial Least Squares Discriminant Analysis (sPLS-DA), coupled with stability and significance tests, was applied for the identification of up or down regulated proteins between breeds. Analyses revealed a total of 25 proteins clearly separating IDU and ILW pigs. Among the top proteins differentiating the two breeds ACAA2 and CES3 were up-regulated in ILW and HIST2H2BF and KHK were up-regulated in IDU. FASN, involved in fatty acid metabolism and encoded by a gene located in a QTL region for fatty acid composition, was up-regulated in ILW. Protein interaction analysis showed that 16 of these proteins were connected in one big module. Functional analyses indicated that differentially expressed proteins were involved in several biological processes related to the metabolism of lipids, amino-acids, carbohydrates, cofactors and antibiotics/drugs, supporting that these functions might distinguish IDU and ILW pigs. This comparative proteomic analysis of the porcine liver highlighted several biological factors that could determine the peculiar production potentials of these two heavy pig breeds, derived by their different genetic backgrounds.

## Introduction

The liver is an important metabolic organ that governs many physiological processes that define biological mechanisms leading to growth, feed efficiency and several other economically relevant traits in all livestock species, including the pig. Metabolism of macronutrients, the capacities to store glucose in the form of glycogen, lipid oxidation and partitioning for deposition in other tissues (e.g. adipose tissues) and catabolism of xenobiotics are among some of the most critical functions of this organ that may contribute to growth rate, feed efficiency and performances of the pigs throughout their lifetime. The liver is also a primary source of proteins and the centre of amino acid metabolism, it is responsible for the secretion of many proteins in the blood and interplays with this circulating tissue for the transfer of amino acids and energy compounds and the disposal of waste metabolites, i.e. derived by the protein degradation in the form of urea metabolism [1, 2].

The construction of liver proteome maps [3, 4] and the establishment of the Human Liver Proteome Project (HLPP) by the Human Proteome Organisation (HUPO) [5] have constituted starting resources for the study of liver function not only in humans but also in all animal species, including the pig [6, 7]. Indeed, a better understanding of the metabolic processes being undertaken in the hepatocytes will be pivotal in managing animal growth and feed efficiency. In this context, proteomics can help to unravel the variations in the metabolic pathways of the hepatocytes in healthy and diseased animals and to identify biomarkers that could be applied in breeding programmes and might be useful to define animal models to better characterize liver physiology and related diseases [8, 9].

Despite these key roles and envisaged potential applications, the interest of the animal science sector on liver proteomics has been slowly increasing over the last years. For example, a few studies have described the dairy cattle liver proteome and compared the liver proteome profiles between different breeds, assuming in this way that differences could be imputed to genetic diversity of the investigated breeds [10]. These studies were applied using traditional proteomic approaches including two-dimensional polyacrylamide gel electrophoresis (2DE) and mass spectrometry (MS) that developed a proteome map of the Holstein liver and compared proteome profiles between a beef cattle breed (i.e. Chianina) and a dairy cattle breed (i.e. Holstein).

A first comprehensive proteomic analysis of porcine hepatic cells was achieved by Caperna *et al.* [11] who analysed three-lines hybrid pigs (Landrace × Yorkshire × Poland China). This study produced 2DE maps of cytosol and membrane fractions from hepatocytes analysed with isoelectric focusing. A total of 728 proteins spots were picked and analysed by MS resulting in a total of 282 unique identified proteins. Wang *et al.* [12] and Liu *et al.* [13] used 2DE and MS to investigate how intrauterine growth restriction affected the liver proteome in new borne and fetal pigs and identified few proteins that differed between the control and treated animals.

More recently, gel-free proteomic approaches have been raised in popularity and have been also applied in livestock. Tang *et al.* [14] investigated the heat stress response in broiler liver using a gel-free method, i.e. sequential window acquisition of all theoretical spectra (SWATH)-MS, reaching a good resolution that was able to quantify about 2400 proteins, one ten of which was differentially expressed between the controls and the heat stressed group. Other MS based proteomics methods, such as label-free liquid chromatography (LC)-MS, have become more popular for quantifying protein expression and compare different samples, giving also the possibility to better investigate hydrophobic proteins with low or high molecular weights that are quite challenging to be analysed using traditional proteomic approaches [15].

It is well known that different pig breeds and lines have different attitudes and potentials for many production and economically relevant traits (like growth rate, feed efficiency, carcass and meat quality traits, reproduction performances and rusticity) that might affect their use or choice in various production systems and breeding programmes. These traits can be considered as external traits or end phenotypes that are the outcome of complex biological processes and interactions that can be dissected only including other intermediate information, which might better describe biological mechanisms underlying their final expression [16]. This intermediate level that links the genetic layer (defined by the genetic variability) and the external phenotypes (i.e. production traits) includes internal phenotypes such as metabolite and proteins quantitative and qualitative profiles [17]. The description of internal phenotypes in pigs that might be easily linked to genetic differences, declined as major differences between pig breeds, has been recently reported comparing the plasma and serum metabolome profiles of two Italian heavy pigs (i.e. Italian Duroc and Italian Large White), commonly used in heavy-pig production systems aimed to produce high quality dry-cured hams [18]. These breeds, that differ for several economically important traits (Italian Large White pigs are mainly selected for meat quality and carcass traits and to maximize maternal reproduction efficiency; Italian Duroc pigs are mainly used as terminal sires and are selected to maximize growth and performance traits and improve meat quality parameters) are commonly included in crossbreeding programmes exploiting heterotic effects to obtain commercial final pigs [19-21].

This study characterized and compared the liver proteomic profiles of Italian Large White and Italian Duroc pigs using a quantitative label-free LC-MS proteomic approach with the final aim to describe intermediate phenotypes that might capture biological differences between these two porcine genetic types and then inform the biological mechanisms underlying their different production performances.

## Materials and methods

All animals used in this study were kept according to the Italian and European legislations for pig production. All procedures described here were in compliance with Italian and European Union regulations for animal care and slaughter. Pigs were not raised or treated in any way for the purpose of this study and for this reason no other ethical statement is needed.

## Animals

Five Italian Duroc gilts and five Italian Large White gilts were included in this study. Animals did not have common grandparents. Gilts from both pig breeds were performance-tested at the Test Station of the National Pig Breeder Association (ANAS). Performance evaluation started when the pigs were 30 to 45 days of age and ended when the animals reached about 155 ± 5 kg live weight. Details of performance testing system in Italian heavy pigs of these two breeds are reported in [22-24].

All pigs were raised, fed and handled in the same ways. After a fasting period of 12 h, all animals were transported in the morning in a commercial abattoir, where were electrically stunned and then slaughtered under controlled conditions. Then, after slaughtering, liver specimens were collected, placed in a labelled tube, snap frozen in liquid nitrogen and then stored at −80 °C until they were analysed.

## Protein extraction and digestion

Protein extraction and digestion were performed according to a slightly modified Filter Aided Sample Preparation (FASP) protocol [25]. Briefly, proteins were extracted from approximately 100 mg of liver tissue homogenised in 1 mL of extraction buffer [0.1% SDS (Affymetrix/Thermo Fisher Scientific, USA), 100 mM Tris/HCl (Affymetrix/Thermo Fisher Scientific, USA) pH 7.6, 10 mM DTT (Affymetrix/Thermo Fisher Scientific, USA)]. Liver samples were homogenised for 2 min using a homogenizer (G50 Tissue Grinder, Coyote Bioscience, Inc., China) in an Eppendorf tube immersed into liquid nitrogen. The lysates were vortexed at room temperature for 3 min and then centrifuged at 16000 × r.c.f. at 4 °C for 5 min. Samples were heated at 56 °C for 30 min and then centrifuged at 16000 × r.c.f. at 20 °C for 20 min. The supernatants were transferred into new labelled tubes, were mixed, aliquoted and stored at −20 °C until usage. Qubit™ Fluorometric Quantitation was used to determine the protein concentration according to the manufacturer’s instructions (Affymetrix/Thermo Fisher Scientific, USA).

Protein samples were reduced and alkylated with DTT and iodoacetamide (IAA, Sigma-Aldrich/Merck, USA) and then digested with trypsin according to the FASP method [25]. Samples were then purified from any contaminants using Pierce C18 Spin Columns (Affymetrix/Thermo Fisher Scientific, USA), vacuum dried and stored at −20 °C. All 10 peptide samples (five from Italian Duroc and five from Italian Large White pigs) were collected for MS analysis.

## Mass spectrometry analysis

Dry peptides from each sample were resuspended in 25 µL of a mixture of water: acetonitrile: formic acid 97:3:2 and sonicated for 10 min at room temperature, then centrifuged at 12,100 × r.c.f. for 10 min. Mass spectrometry analyses were performed on an ESI-Q-TOF Accurate-Mass spectrometer (G6520A, Agilent Technologies, Santa Clara, CA, USA), controlled by MassHunter software, Agilent (v. B.04.00) (https://www.agilent.com/en/products/software-informatics/masshunter-suite/masshunter/masshunter-software) and interfaced with a CHIP-cube to an Agilent 1200 nano-pump (Agilent Technologies, Santa Clara, CA, USA).

Chromatographic separation was performed on a high-capacity loading chip (Agilent Technologies) with a 75 µm internal diameter (I.D.), 150 mm, 300 Å C18 column, prior to a desalting step through a 500 nL trap column. The injected samples (8 μL) were loaded onto the trap column with a 4 μL/min 0.1% (v/v) formic acid (FA):acetonitrile (ACN) (98:2 v/v) phase flow. After 3 min, the precolumn was switched in-line with the nanoflow pump (400 nL/min, phase A: water:ACN:FA (96.9:3:0.1 v/v/v), phase B: ACN:water:FA 94.5:5:0.1 v/v/v), equilibrated in 1% (v/v) B. The peptides were eluted from the reverse phase (RP) column through the following gradient: 1% for one min, 1->5% B in 7 min, 5->30% B over a period of 142 min, 30–60% B in 20 min, 60->90% B in 0.1 min, then held at 90% B for 8 min, and switched back to 1% B for column reconditioning, for a total runtime of 190 min. Eight µL of each samples were run twice; analytical controls (a mix of baker’s yeast enolase and bovine serum albumin tryptic digests) were run daily to monitor chromatographic performances. Ions were formed in a nano-ESI source, operated in positive mode, 1860 V capillary voltage, with the source gas heated at 350 °C and at a 5 L/min flow. Fragmentor was set to 160 V, skimmer lens operated at 65 V. Centroided MS and MS^2^ spectra were recorded from 250 to 1700 m/z and 70 to 1700 m/z, respectively, at scan rates of 8 and 3 Hz. The six most intense multi-charged ions were selected for MS^2^ nitrogen-promoted collision-induced dissociation. The collision energy was calculated according to the following expression: 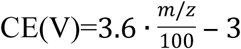. A precursor active exclusion of 0.22 min was set, and the detector was operated at 2 GHz in extended dynamic range mode. Mass spectra were automatically recalibrated with two reference mass ions.

## Label-free quantitative profiling

Raw MS data were converted as MASCOT (www.matrixscience.com) generic file (mgf) using MassHunter Qualitative Analysis (v. B.05.00, Agilent Technologies) and used for peptides identification with MASCOT (version 2.4, Matrix Science, London, UK) searched against an implemented version of the Pig PeptideAtlas resource (http://www.peptideatlas.org/) [26], including contaminant protein sequences as retrieved from the cRAP v.1.0 resource (http://www.thegpm.org/crap/). The search parameters used were as follow: 40 ppm precursor tolerance, 0.1 Da fragment mass error allowed, two missed cleavage allowed for trypsin, carbamidomethyl as a fixed modifier of cysteine residues, asparagine and glutamine deamidation and methionine oxidation as variable modification. The peptide identification results were imported into an in-house Trans-Proteomic Pipeline server (TPP, v. 5.1.0;) [27], rescored and validated with PeptideProphet [28]. Protein inference was then achieved by ProteinProphet [29]. As last step, relative label-free quantification was performed with Skyline v.4.1 [30] using the rescored search results (Protein probability ≥0.9, corresponding to a global FDR≤0.1). PepXML files and raw mzXML data were imported in Skyline and: (i) the protein database fasta were filtered by retaining only the proteins with a ProteinProphet score ≥ 0.9 and (ii) MS^1^ data were correlated to the corresponding peptide identification, matches were filtered (idotP≥0.9). Data were exported and analyzed as following described.

Data were imported and analyzed with InfernoRDN v.1.1.6556.25534 (https://omics.pnl.gov/software/InfernoRDN; [31]). Data were Log2 transformed and normalized with the central tendency adjustment method. Peptide measurements were then rolled up to corresponding protein abundances through the Zrollup method. In the ZRollup procedure a scaling method similar to z-scores is applied first to peptides that originate from a single protein and then the scaled peptide measures are averaged to obtain a relative protein abundance measure [31]. Differences in protein abundance between Italian Large White and Italian Duroc pigs were investigated as described below. Proteins were considered only if identified with more than one peptide.

The mass spectrometry proteomics data have been deposited to the ProteomeXchange Consortium via the PRIDE (http://www.ebi.ac.uk/pride; [32]) partner repository with the dataset identifier PXD009771 and project DOI: 10.6019/PXD009771.

## Data analyses

### Multivariate statistical analysis

Differentially abundant proteins were detected by applying the multivariate approach of sparse Partial Least Squares Discriminant Analysis (sPLS-DA) [33] coupled with the validation procedure detailed in Bovo *et al.* [18, 34]. sPLS-DA is a multivariate technique used in classification and discrimination problems especially when variables are highly correlated [35]. Briefly, breed was modeled as response variable (Italian Duroc = 0, Italian Large White = 1) and proteins as predictors. The sPLS-DA penalization coefficient *eta* (ranging from 0.1 to 0.9) and the number of hidden components *K* (ranging from 1 to 5) were automatically selected by an internal 4-fold cross-validation procedure (4CV). The selected proteins were then assessed through a stability and significance test [18]. The stability test was based on a Leave One Out (LOO) procedure coupled with a permutation test. For this purpose, 1,000 artificial datasets were obtained by randomly permuting the breed trait values. While the stability test is aimed at evaluating the frequency of selection of a protein in the original dataset against the permuted ones, the significance test evaluates the regression coefficient (β). These two tests estimate the probability (*P*) that the selection of a given protein inside the dataset is due to chance or to a particular structure of the dataset. Proteins having a *P* < 0.10 (with the sign of the regression coefficient that matched the fold change ratio) were declared stable and significantly related to breed. Analyses were performed in R v. 3.0.2 [36] by using the “spls” package (function “cv.splsda” and “splsda”). Scatter plot of the first two components was drawn (each point represents an individual sample).

### Functional and protein network analyses and relation to genomic information

Functional interpretation of differentially abundant proteins was carried out in Cytoscape (http://www.cytoscape.org/) [37] using the plug-in ClueGO (http://www.ici.upmc.fr/cluego/) [38]. Gene enrichment analysis was carried out over the Gene Ontology (GO) – Biological Process (BP) branch (release data: May 2018) by setting the following parameters: (i) GO hierarchy level from 3 to 20; minimum number of genes per GO term equal to 2; minimum of input genes associated to the functional term greater than 2%; GO term network connectivity (Kappa score) equal to 0.70 and right-sided hypergeometric test. This analysis made use of *Sus scrofa* specific functional annotations. The other parameters were kept with default values. GO:BP terms with a Benjamini–Hochberg corrected p-value < 0.05 were considered statistically over-represented. A pathway enrichment analysis was separately carried out, with ClueGO, over the KEGG pathway database (release data: May 2018).

Protein-protein interaction (PPI) analysis of differentially abundant proteins identified in the breed comparison was obtained using STRING v. 10.5 database (https://string-db.org/) [39]. STRING is a web resource providing uniquely comprehensive coverage of experimental and predicted interaction information. The analysis was carried out considering the *Sus scrofa* specific interactome. Only interactions having a STRING combined score > 0.4 were considered (i.e medium confidence). Network indices such as the number of nodes and edges, the average node degree (average no. of connections), the expected number of edges and the PPI enrichment *p*-value were computed. The expected number of edges gives how many edges is to be expected if the nodes were to be selected at random. A small PPI enrichment *p*-value indicates that nodes are not random and that the observed number of edges is significant (https://string-db.org/).

The Pig Quantitative Trait Locus Database (Pig QTLdb release 35) [40] was used to explore possible links between up or down-regulated proteins and genomic QTLs for economically relevant traits and parameters that could be affected by genes encoding the these proteins. The whole set of 27,465 QTLs/associations was downloaded and their genome coordinates where compared with genome coordinates of the genes encoding differentially abundant proteins observed in this study. Borders of the considered genomic regions were defined by the gene coordinates ± 50 kbp. QTL relationships were filtered by manually curating functional relationships of the genes encoding these proteins.

## Results

### Label-free proteomic data of porcine liver samples

Label-free proteomic analysis was performed from proteins extracted from the liver of pigs of two heavy pig breeds, Italian Duroc and Italian Large White. The analysis of the Italian Duroc liver specimens identified a total of 1,696 peptides belonging to 467 proteins, with an average of 3.6 peptides per protein. The analysis in the Italian Large White liver samples identified 1,253 peptides related to 350 proteins, with an average of 3.6 peptides per protein. Combining these two datasets, this study identified a total of 1,873 peptides related to 501 proteins (average of 3.7 peptides per protein), that were then ri-defined using stringent parameters in the InfernoRDN analysis, obtaining 200 unique proteins (including a total of 1,041 peptides; S1 Table). These proteins were included in a total of 13 GO:Biological Processes (Fig. 1), four of which (organonitrogen compound metabolic process, lipid metabolic process, carbohydrate metabolic process and cellular amino acid metabolic process) accounted for more than 10% of the total listed proteins.

Only proteins having at least two supporting peptides were kept and subsequently used in the study. Bioinformatics analyses were carried out on a final dataset counting 200 proteins (including a total of 1,041 peptides) identified with high confidence and subsequently quantified. The full list of the 200 proteins analysed in this study, as well as the information on the peptides identified, is shown in the S1 Table.

**Fig. 1.**
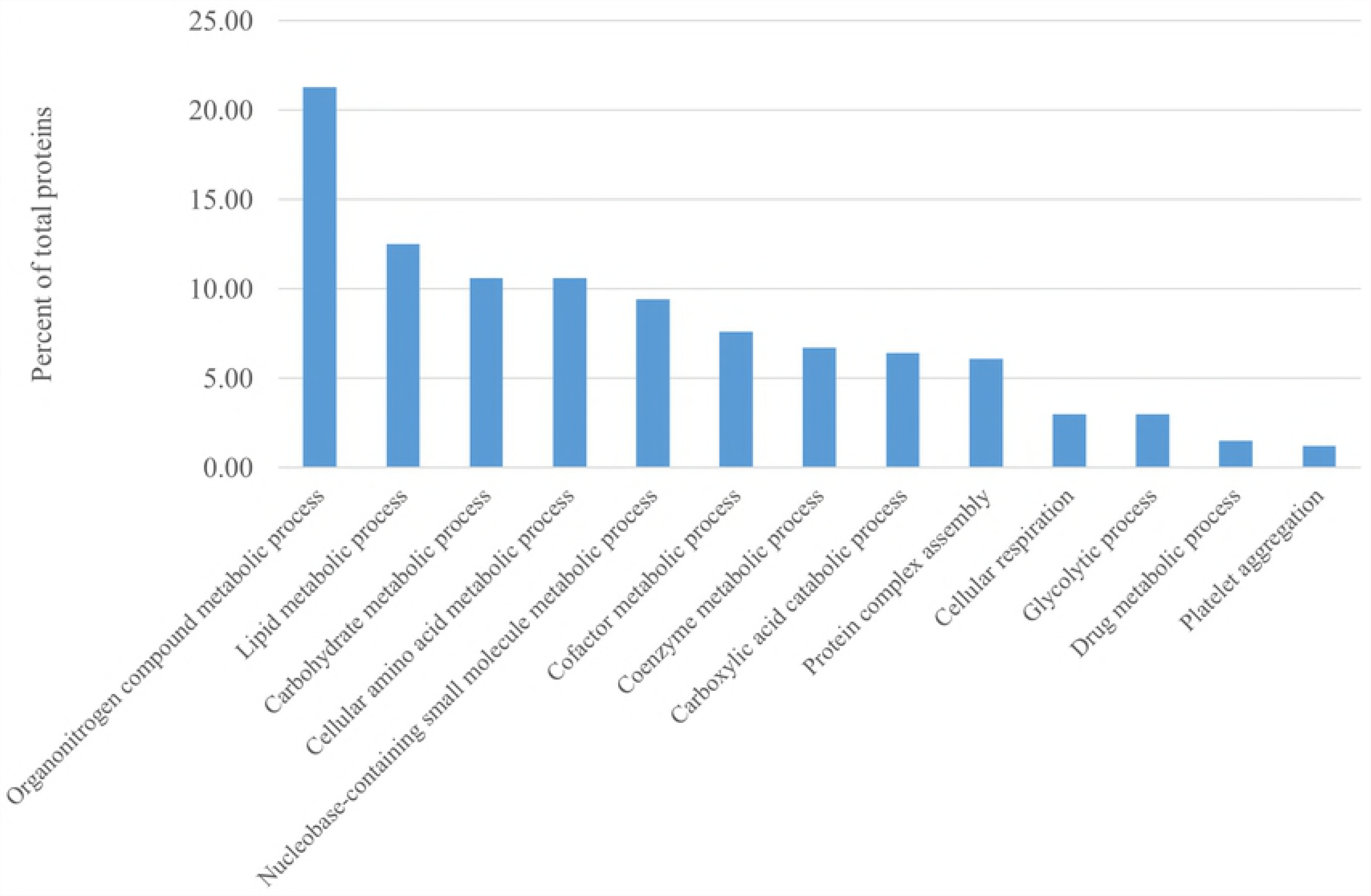
Percentage of liver proteins (over all 329 identified proteins; S2 Table) grouped according to different biological processes.

### Comparative analysis of breed derived proteomic liver profiles

Differences at the proteome level between Italian Large White and Italian Duroc pigs were investigated by using the multivariate approach of sPLS-DA. This technique was coupled with a statistical procedure aimed at evaluating the stability and significance of the proteins selected as differentially expressed. The scatter plot of the first two sPLS-DA components shows that animals of the same breed clustered together, indicating that the identified proteins can discriminate these two heavy pig populations (Fig. 2). According to the stability test, 33 proteins (16.5% on the total) had a *P* ≤ 0.10 (defined as a threshold of significance, based on the validation procedure; [18]). The sPLS-DA regression coefficient of 57 proteins (28.5%) was equal to 0, indicating that these proteins do not have any weight in the classification derived by the breed. At the significance test, 56 out of 57 proteins had a *P* ≤ 0.10. A total of 25 proteins out 200 identified proteins (12.5%) resulted both stable (in terms of statistically assigned condition) and significantly differentially expressed (*P* ≤ 0.10) (Table 1). Among these proteins, 14 (56%) showed an increased abundance in Italian Duroc pigs, while the remaining 11 (44%) had a higher quantification in Italian Large White pigs. Details about proteins with significant different quantification between the two breeds are presented in Table 1. Four of these proteins (two with the highest expression in Italian Duroc pigs: 3-ketoacyl-CoA thiolase, mitochondrial (ACAA2) and histone H2B type 2-F (HIST2H2BF); and two with the highest expression in Italian Large White pigs: ketohexokinase (KHK) and carboxylesterase 3 (CES3)) showing one of the two calculated *P* values ≤0.01 and the other with ≤0.05, were considered the top proteins identified in this study differentiating the two analysed breeds.

**Fig. 2.**
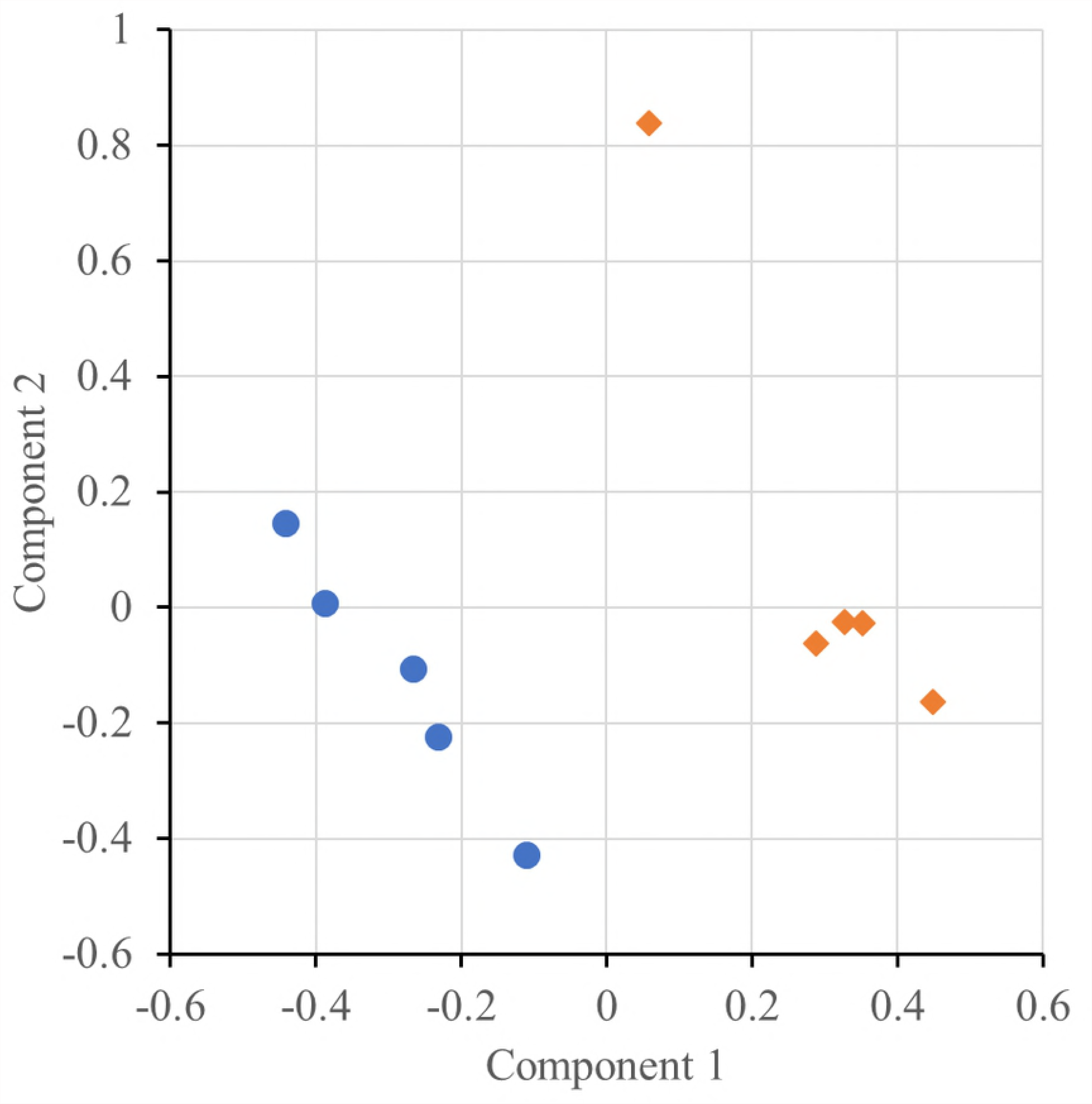
Scatter plot of the first two sPLS-DA components obtained from the liver proteome of five Italian Duroc pigs (blue circles) and five Italian Large White pigs (orange diamonds).

**Table 1.**
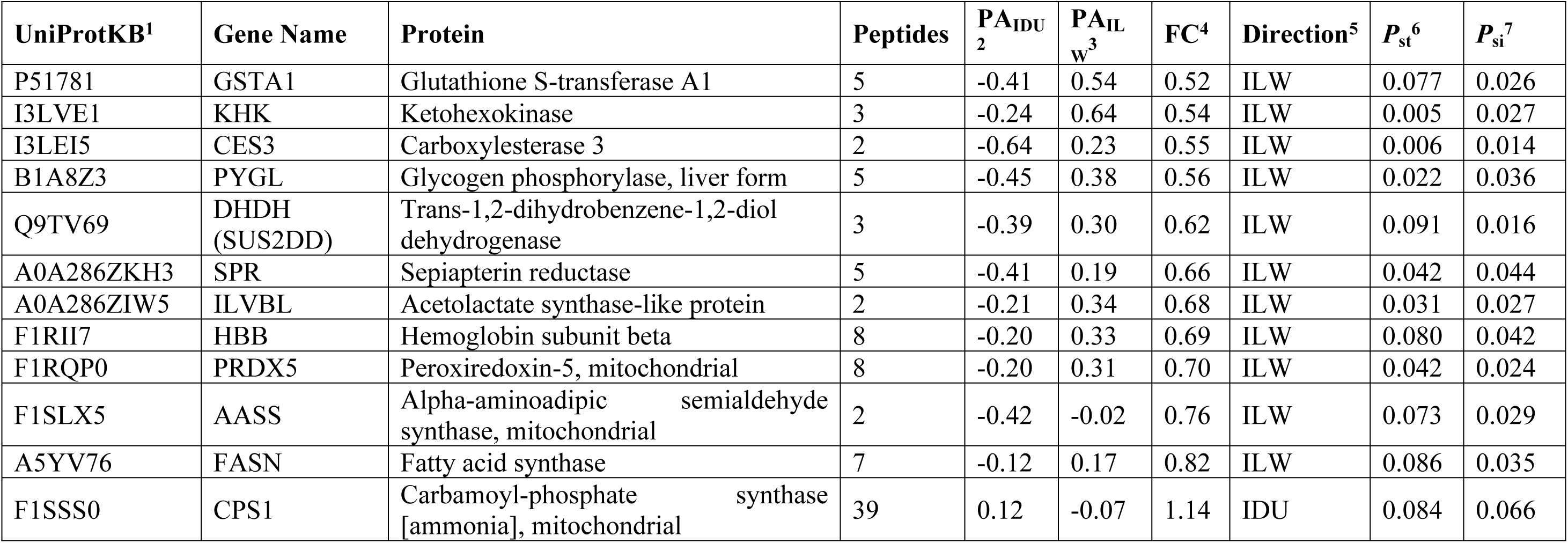
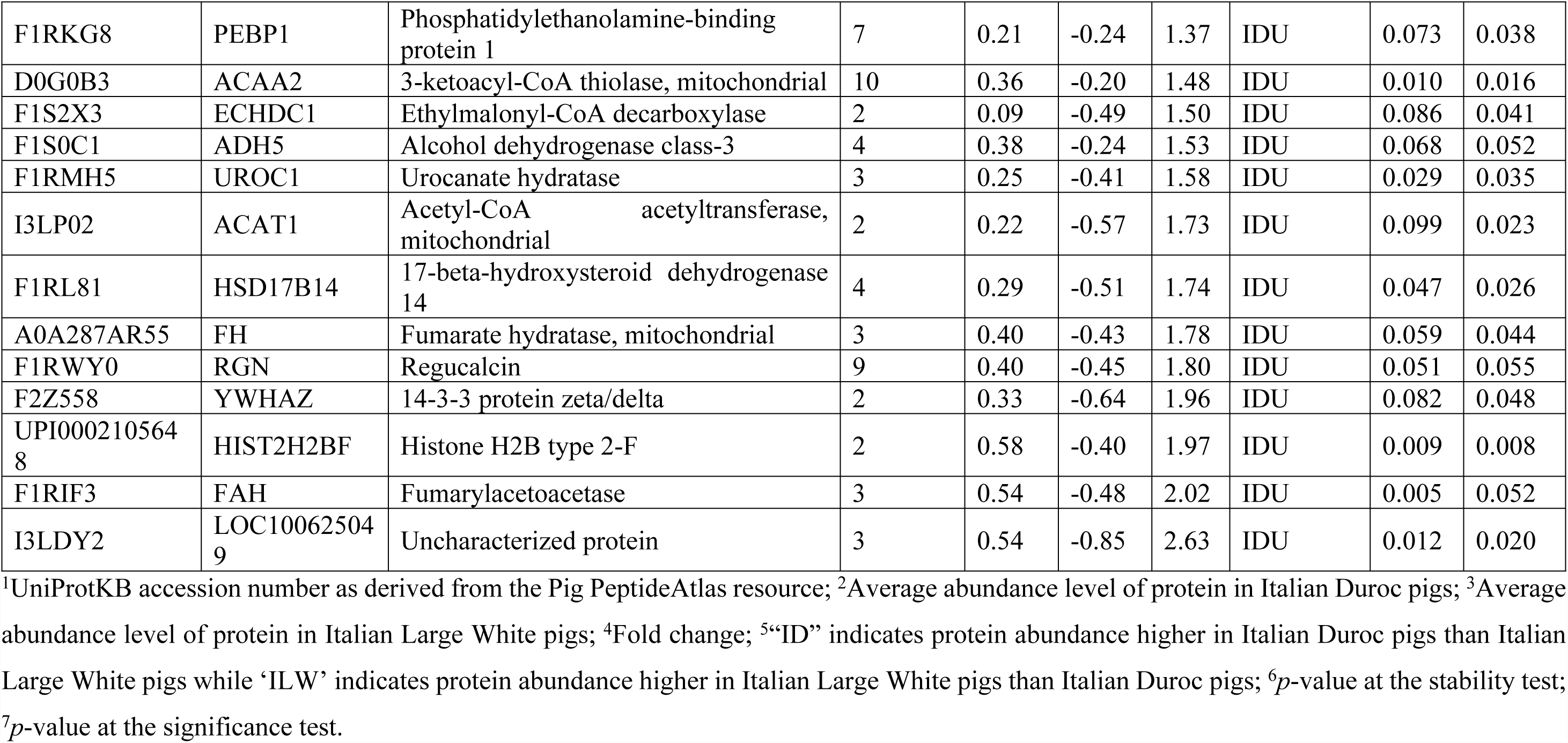
Differentially abundant proteins identified from label-free mass spectrometry analysis of liver samples of Italian Duroc (ID) and Italian Large White (ILW) pigs.

### Functional inference of proteomic differences between breeds

Biological processes and pathways involving the 25 differentially abundant proteins were investigated through functional association analysis. Enrichment analyses were carried out in Cityscape (ClueGO package) by setting the significance threshold at 0.05 on the Benjamini-Hotchberg corrected p-values. Analyses run over the GO:BP and KEGG pathway databases, separately. Two proteins (HIST2H2BF and LOC100625049) were not in the ClueGO annotation sets. A total of 24 GO:BP terms (involving 14 differentially abundant proteins) were retrieved (Table 2, Fig. 3A). These biological processes can be summarized as follows: (i) metabolism of lipids, (ii) metabolism of amino-acids, (iii) metabolism of carbohydrates, (iv) metabolism of cofactors and (v) metabolism of antibiotics/drugs. It is interesting to note that processes related to the glutamine family amino acids and lipids involve proteins with a higher abundance in Italian Duroc than in Italian Large White pigs. Processes related to organic acid catabolism cellular/alpha amino acid metabolism showed the same direction, except for the Aminoadipate-Semialdehyde Synthase (AASS) protein, that was more expressed in the Italian Large White pigs. This inversion was also evident for proteins involved in the carbohydrate catabolic process that showed a higher abundance in Italian Large White than in Italian Duroc pigs.

**Table 2.** Over-represented biological processes (GO:BP) associated to up or down regulated proteins in the breed comparison.

| Term | Description | p-value<sup>1</sup> | % of associated proteins<sup>2</sup> | N. of proteins | Up or down regulated proteins<sup>3</sup> |
| --- | --- | --- | --- | --- | --- |
| GO:0005996 | monosaccharide metabolic process | 2.26E-03 | 2.05 | 3 | KHK↓, RGN↑, SUS2DD↓ |
| GO:0006091 | generation of precursor metabolites and energy | 1.23E-04 | 2.03 | 5 | AASS↓, ACAT1↑, ADH5↑, FH↑, PYGL↓ |
| GO:0016052 | carbohydrate catabolic process | 7.63E-03 | 2.47 | 2 | PYGL↓, SUS2DD↓ |
| GO:0016999 | antibiotic metabolic process | 6.36E-03 | 2.78 | 2 | ADH5↑, FH↑ |
| GO:0042737 | drug catabolic process | 5.31E-03 | 3.13 | 2 | AASS↓, ACAT1↑ |
| GO:0009108 | coenzyme biosynthetic process | 1.40E-03 | 2.48 | 3 | ACAT1↑, RGN↓, SPR↓ |
| GO:0008652 | cellular amino acid biosynthetic process | 2.95E-03 | 4.44 | 2 | AASS↓, CPS1↑ |
| GO:1901607 | alpha-amino acid biosynthetic process | 2.86E-03 | 4.65 | 2 | AASS↓, CPS1↑ |
| GO:0009064 | glutamine family amino acid metabolic process | 3.05E-03 | 4.26 | 2 | CPS1↑, FAH↑ |
| GO:0006525 | arginine metabolic process | 2.81E-04 | 16.67 | 2 | CPS1↑, FAH↑ |
| GO:0016042 | lipid catabolic process | 4.45E-05 | 2.59 | 5 | ACAA2↑, ACAT1↑, CPS1↑, ECHDC1↑, HSD17B14↑ |
| GO:0044242 | cellular lipid catabolic process | 1.16E-04 | 3.25 | 4 | ACAA2↑, ACAT1↑, CPS1↑, ECHDC1↑ |
| GO:0034440 | lipid oxidation | 2.82E-04 | 4.62 | 3 | ACAA2↑, ACAT1↑, ECHDC1↑ |
| GO:0019395 | fatty acid oxidation | 2.81E-04 | 4.76 | 3 | ACAA2↑, ACAT1↑, ECHDC1↑ |
| GO:0009062 | fatty acid catabolic process | 2.54E-04 | 5.08 | 3 | ACAA2↑, ACAT1↑, ECHDC1↑ |
| GO:0072329 | monocarboxylic acid catabolic process | 2.89E-04 | 4.35 | 3 | ACAA2↑, ACAT1↑, ECHDC1↑ |
| GO:0006635 | fatty acid beta-oxidation | 1.33E-04 | 6.52 | 3 | ACAA2↑, ACAT1↑, ECHDC1↑ |
| GO:0044282 | small molecule catabolic process | 2.21E-07 | 3.85 | 7 | AASS↓, ACAA2↑, ACAT1↑, ECHDC1↑, FAH↑, SUS2DD↓, UROC1↑ |
| GO:0016054 | organic acid catabolic process | 5.77E-07 | 4.51 | 6 | AASS↓, ACAA2↑, ACAT1↑, ECHDC1↑, FAH↑, UROC1↑ |
| GO:0006520 | cellular amino acid metabolic process | 4.45E-05 | 2.59 | 5 | AASS↓, ACAT1↑, CPS1↑, FAH↑, UROC1↑ |
| GO:1901605 | alpha-amino acid metabolic process | 1.00E-05 | 3.82 | 5 | AASS↓, ACAT1↑, CPS1↑, FAH↑, UROC1↑ |
| GO:0009063 | cellular amino acid catabolic process | 8.06E-06 | 7.14 | 4 | AASS↓, ACAT1↑, FAH↑, UROC1↑ |
| GO:0046395 | carboxylic acid catabolic process | 5.77E-07 | 4.51 | 6 | AASS↓, ACAA2↑, ACAT1↑, ECHDC1↑, FAH↑, UROC1↑ |
| GO:1901606 | alpha-amino acid catabolic process | 7.81E-06 | 8.16 | 4 | AASS↓, ACAT1↑, FAH↑, UROC1↑ |

<sup>1</sup>Benjamini-Hochberg corrected *p*-values; <sup>2</sup>Percentage of input proteins found associated with respect to the number of proteins directly annotated with the functional term; <sup>3</sup>The symbol ↑ indicates protein abundance higher in Italian Duroc pigs than Italian Large White pigs while the symbol ↓ indicates protein abundance higher in Italian Large White pigs than Italian Duroc pigs

**Fig. 3.**
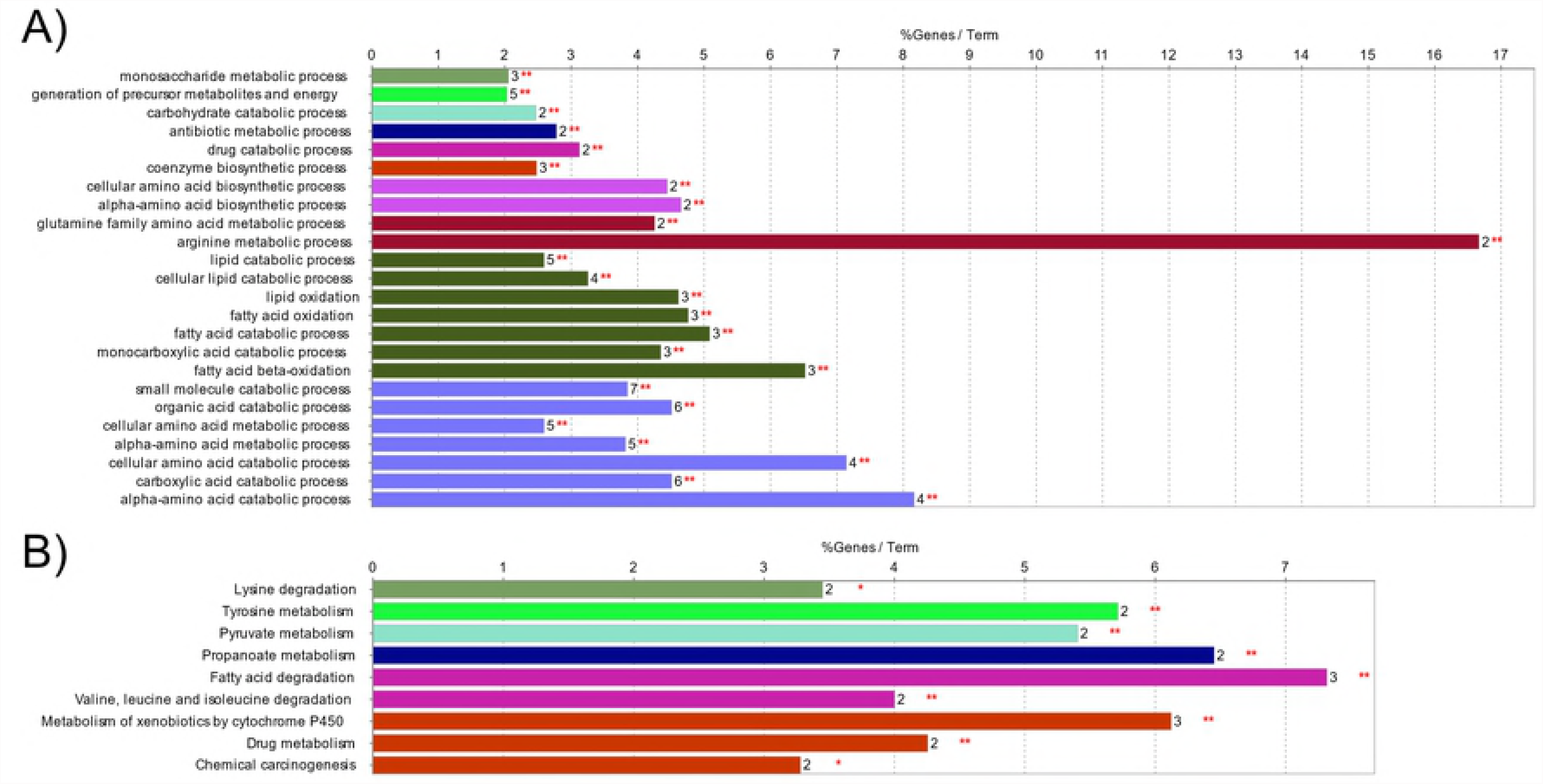
Functional analysis of the 25 differentially abundant proteins between Italian Duroc and Italian Large White pigs. The two panels show the result of gene enrichment analyses over the A) Gene Ontology – Biological Process branch, and B) over the KEGG pathway database. Bars represent the percentage of input proteins found associated with respect to the number of proteins directly annotated with the functional term. The number of input proteins related to the term and the term significance are reported next to each bar. Detailed statistics are reported in Table 2 and Table 3, respectively. In each panel, bars sharing a specific color are clustered in the same functional group.

Over-representation analysis over the KEGG pathway database highlighted a total of 9 pathways (Table 3, Fig. 3B) related to the metabolism of lipids, amino-acids, carbohydrates and chemicals (as previously showed from the analysis over the GO:BP database). Nine proteins were involved in these pathways. Again, in this case, all the differentially abundant proteins involved in the fatty acid catabolism had higher concentration in Italian Duroc pigs than in Italian Large White pigs. Moreover, in addition to the glutamine family amino acids, the pathways involving valine, leucine, isoleucine and tyrosine showed the same behaviour.

**Table 3.** Over-represented KEGG pathways associated to up or down regulated proteins in the breed comparison.

| Term | Description | p-value<sup>1</sup> | % of associated proteins<sup>2</sup> | N. of proteins | Up or down regulated proteins<sup>3</sup> |
| --- | --- | --- | --- | --- | --- |
| KEGG:00310 | Lysine degradation | 1.06E-02 | 3.45 | 2 | AASS↓, ACAT1↑ |
| KEGG:00350 | Tyrosine metabolism | 7.88E-03 | 5.71 | 2 | ADH5↑, FAH↑ |
| KEGG:00620 | Pyruvate metabolism | 7.04E-03 | 5.41 | 2 | ACAT1↑, FH↑ |
| KEGG:00640 | Propanoate metabolism | 8.27E-03 | 6.45 | 2 | ACAT1↑, ECHDC1↑ |
| KEGG:00071 | Fatty acid degradation | 1.28E-03 | 7.32 | 3 | ACAA2↑, ACAT1↑, ADH5↑ |
| KEGG:00280 | Valine, leucine and isoleucine degradation | 9.07E-03 | 4.00 | 2 | ACAA2↑, ACAT1↑ |
| KEGG:00980 | Metabolism of xenobiotics by cytochrome P450 | 1.09E-03 | 6.12 | 3 | ADH5↑, GSTA1↓, SUS2DD↓ |
| KEGG:00982 | Drug metabolism | 9.38E-03 | 4.26 | 2 | ADH5↑, GSTA1↓ |
| KEGG:05204 | Chemical carcinogenesis | 1.04E-02 | 3.28 | 2 | ADH5↑, GSTA1↓ |

<sup>1</sup>Benjamini-Hochberg corrected *p*-values; <sup>2</sup>Percentage of input proteins found associated with respect to the number of proteins directly annotated with the functional term; <sup>3</sup>The symbol ↑ indicates protein abundance higher in Italian Duroc pigs than Italian Large White pigs while the symbol ↓ indicates protein abundance higher in Italian Large White pigs than Italian Duroc pigs.

STRING was used to highlight the functional connections established among the differentially abundant proteins. The analysis revealed a connected protein network (Fig. 4) divided in: (i) one big module composed by 16 nodes (64%) and 19 links, (ii) a small component of two proteins (8%) and (iii) seven singletons (28%). The resulting network showed a PPI enrichment *p*-value of 3.67×10^−10^ (three expected edges vs. 20 detected edges) indicating that proteins are at least partially biologically connected. In this network most of the proteins interacted with only one or two other partners (average node degree equal to 1.6). However, two proteins (ADH5 and HSD17B14; with higher expression in Italian Duroc than in Italian Large White pigs) presented the highest degree of connection (six edges), which suggest them as “hub” proteins playing a putative role of controllers inside biochemical pathways that could potentially lead to cascade of protein expression differences. In addition, the big module included all proteins that were included in the GO and KEGG enriched processes that clearly differentiated (in terms of direction of the relative level of expression) Italian Duroc and Italian Large White liver proteomic profiles.

**Fig. 4.**
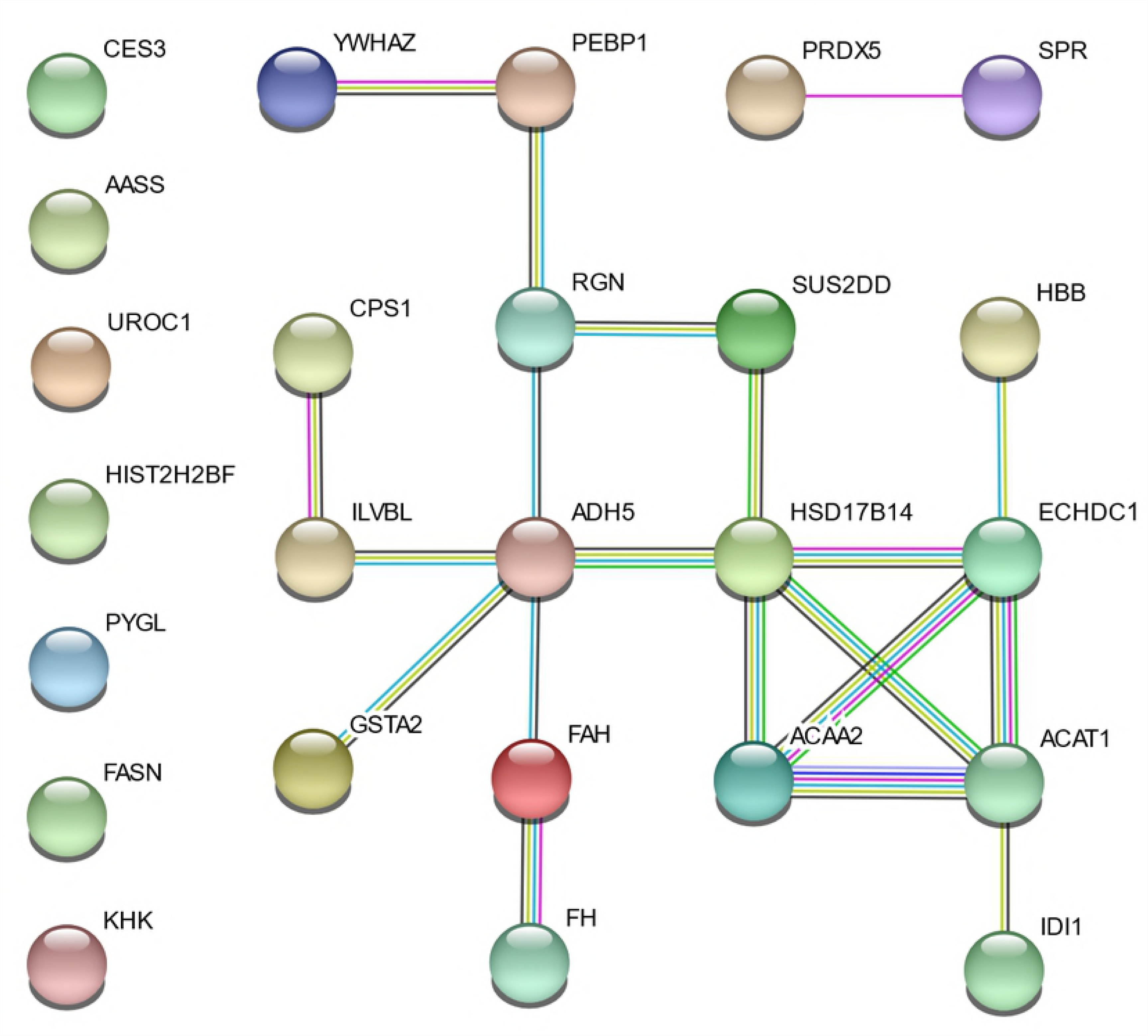
Protein-protein interaction map of the 25 differentially abundant proteins. Interactions are based on STRING v.10.5 and are shown in different colors: cyan is from curated databases, magenta is experimentally determined, dark green is gene neighborhood, blue is gene co-occurrence, light green is textmining, black is co-expression and purple is protein homology.

S3 Table reports information on the potential links between differentially expressed proteins identified in the comparison between breeds and QTL mapped in the corresponding gene regions, as retrieved from the pigQTL database. A total of twelve out of 25 up or down regulated proteins identified in the breed comparative analysis resulted located in ± 50 kbp surrounding 23 QTL regions (for a total of 40 protein/QTL combinations), that might be directly or indirectly (in a broad sense) affected by the protein functions.

Six of these proteins were up-regulated in Italian Duroc pigs (ECHDC1, HSD17B14, FAH, UROC1, ACAT1, CPS1) and six in Italian Large White pigs (PRDX5, SPR, GSTA1, FASN, KHK, CES3).

## Discussion

Despite the fact that the pig is one of the most important animal species used as protein source for human consumption and that is also considered a suitable model for several biomedical aspects related to liver functions [41, 42], few studies have investigated the pig liver proteome compared to what is available for other species [9, 11-13, 43-52]. Most of these liver proteomic investigations compared proteomic profiles of pigs under different experimental conditions or treatments or just wanted to extend the porcine liver proteome map. Our work not only contributed to improve a more detailed picture of the protein expressed in pig liver but also provided a first comparative analysis of the liver proteome of two important heavy pig breeds, Italian Duroc and Italian Large White. These breeds are mainly used in crossbreeding programmes to obtain commercial slaughtered pigs. As the liver can be considered a key organ for translating growth rate, feed efficiency and performance potentials of the animals by exploiting tissue specific metabolic functions, the evaluation of proteome differences between these Italian Duroc and Italian Large White could highlight biological aspects (that, in turn, are derived by genetic factors) that distinguish and characterize these breeds and that might determine their peculiar production potentials.

It is clear that any proteomic study cannot disclose in a single analysis or method all proteins that construct a tissue or an organ for intrinsic limits of the available technologies and difficulties to discriminate and separate all proteins present in complex matrix. The predicted human proteome has been estimated to be constituted by at least 20,000 proteins (without considering splicing variants), about 59% of which might be expressed in the liver [53]; a similar fraction could be expected in the porcine liver. Therefore, the liver could be considered one of the richest organs in terms of number of proteins. However, studies in pig liver reported, in most cases, tens or just a few hundreds of proteins. For example, Caperna *et al.* [11] using 2DE produced two liver porcine proteomic maps, one from the cytosol fraction and another one from the membrane fraction. Following MS analysis, a total 282 proteins were identified. Golovan *et al.* [44] using isobaric tag for relative and absolute quantification (iTRAQ) based proteomics identified a higher number of proteins (i.e. n. = 880) in the pig liver proteomic profiles obtained from two pig lines (transgenic Enviropig and conventional Yorkshire).

In our study, the label-free LC-MS analysis revealed a total of 501 different proteins that, after stringent filtering were identified as 329 unique proteins, of which 200 were then considered in the comparison between the two breeds (S1 and S2 Tables). Identified proteins overlapped, at least in part, the proteomic general profiles of previous studies carried out in pigs or other livestock species [11, 44, 54]. Most of the identified proteins were involved in several metabolic processes that characterize the liver functions (Fig. 1). Differences among studies might be due to different methods of protein extraction employed and by the peptide separation and identification technologies that could be more or less efficient, also considering the throughput and resolution potentials that determine, at the end, the number of detected proteins. For example, the 2DE study of Caperna *et al.* [11] identified a total of 282 unique porcine liver proteins, the majority of which were of the cytosol fractions. It is well know that this method is technically challenging and is limited for the analysis of very hydrophobic and/or membrane proteins [55]. Using iTRAQ, Golovan *et al.* [44] identified a higher range of protein in the pig proteome, but only few of them (i.e. two and four) were differentially expressed between breeds and sexes, respectively. In this method (label-based), stable isotopes are employed to create specific mass tags that can be recognized by the mass spectrometer and to provide, at the same time, the basis for quantification. One of the main advantage of this method is that allows, in a single MS run, the simultaneous analysis of many samples, reducing analytical variability [56]. On the other hand, the method that we applied (label free approach) provides a higher dynamic range of quantification than stable isotope labelling methods [57].

To our knowledge, this work is the first study applying a label-free method to unravel the pig liver proteome. In the label-free study of Miller *et al.* [54], that compared the liver proteome of two sheep breeds, a larger number of proteins was identified (2,445 vs 501 of our work), probably due to differences in sensitivity of the used instruments. Miller *et al.* [54] and our work had a similar objective, i.e. identify proteomic differences between breeds. In both studies that were based on a similar experimental design (both included comparative analyses between breeds), the same percentage of proteins (about 8%) were declared as differentially expressed between the two analysed genetic types (within species). Even if it could be quite speculative, this general overlapping gross picture might define similar levels of underlying genetic diversity between the compared groups of the two species.

As resulted from the obtained sPLS-DA plot, pigs of the two breeds that we investigated here are well separated by their liver proteomic profiles (Fig. 2). This result is almost overlapping to the results we obtained analyzing targeted metabolomic profiles of plasma and serum of Italian Duroc and Italian Large White pigs [18]. Liver interplays with blood exchanging the products of their respective anabolism and catabolism and many other components, depicting a metabolic picture that could be observed at the organ or tissue level as well in the circulating tissue and in its fractions, i.e. plasma and serum.

Among the 25 proteins that were up-regulated in one breed and, on the other hand, down-regulated in the other breed we could mention a few of them that, according to their function, might highlight some of the differences between Italian Duroc and Italian large White pigs and, in part, explain some of the metabolomic profiles already evidenced [18]. Carboxylesterase 3 (CES3), one of the top significant proteins, was up-regulated in Italian Large White. This is a member of enzymes of the endoplasmic reticulum that hydrolyses a variety of esters, carbamates, amides and similar structures of drugs and xenobiotics [58]. This enzyme is involved in hepatic very low-density lipoprotein (VLDL) assembly and in basal lipolysis of adipose tissues. Ablation of carboxylesterase 3 expression in mice results in reduced circulating plasma triacylglycerol, apolipoprotein B, and fatty acid levels and increased food intake and energy expenditure [59, 60]. Moreover, the porcine CES3 gene is located on SSC6 in a QTL region for residual feed intake (that might be related to feed intake and metabolism efficiency). Differences of hepatic CES3 levels between Italian Large White and Italian Duroc might be in line with a higher feed efficiency of latter breed. ACAA2, among the top up-regulated proteins in Italian Duroc pigs, is a mitochondrial enzyme (3-ketoacyl-CoA thiolase, mitochondrial) catalysing the last step in fatty acid oxidation to release acetyl CoA for the cycle of Krebs. For its role, it is considered one of the key enzymes in lipid metabolism [61], further evidencing differences related to fat related pathways. Again, ACAT1, that is involved in cellular cholesterol homeostasis, macrophage cholesterol metabolism, isoleucine metabolism and ketogenesis (i.e. Hai *et al.* [62]), was up-regulated in Italian Duroc pigs. The *ACAT1* gene is located on porcine chromosome (SSC) 9 in a region in which several QTLs for average daily gain and intramuscular fat content have been identified (S3 Table). In addition, this protein has been already identified to be differentially expressed in a proteomic analysis of skeletal muscles between pigs with extreme values of intramuscular fat content [63], strengthening a role of lipid metabolism differences between pig breeds that could result in different phenotypic traits [23, 64]. Moreover, a few other hepatic differences between Italian Duroc and Italian Large White pigs highlighted also metabolic differences in fatty acid metabolism. The *ECHDC1* gene, which encodes for a protein involved in the mitochondrial fatty acid oxidation, is located in a region of the SSC1 overlapping a QTL region involved in the saturated fatty acid content (S3 Table). FASN (up-regulated in Italian Large White pigs) is an enzyme that catalyses the biosynthesis of palmitic acid. The porcine *FASN* gene is located on SSC12 in a QTL region in the region in which two QTLs affecting the myristic and palmitic acids, have been mapped and for which *FASN* was pointed out as the most plausible candidate gene [65]. Histone H2B (HIST1H2BA), is a member of the histones family, which are basic nuclear proteins that are responsible for the nucleosome structure of the chromosomal fiber in eukaryotes [66]. This family of proteins shown antimicrobial properties in many species [67-69]. For example, Li et al [68] highlighted the antimicrobial activity of HIST1H2BA and other histone proteins in the chicken liver extract. In our study, HIST2H2BF was up-regulated in the Italian Duroc breed, which may allude to a stronger resistance to disease of this breed, that is, in general considered the most rustic among all commercial pig breeds [70].

These specific differences highlighted by the function of the proteins mentioned above (and all other proteins that contributed to produce “breed specific” proteomic profiles; Table 1) can be summarized in few biological processes that could be identified summing up their roles, matching again some of the metabolomic differences (described with biogenic amines and sphingomyelins) that we already reported between these two breeds [18]. These biological processes can be mainly described with metabolism of lipids, metabolism of amino-acids, metabolism of carbohydrates, metabolism of cofactors and metabolism of antibiotics/drugs that, on the whole, embrace all major hepatic functions. Liver proteomic profiles we obtained for Italian Duroc and Italian Large White pigs seems to match production characteristics of these breeds that have been developed over decades of divergent directional selection originally determined by the genetic pools constituting these two heavy pig breeds.

## Conclusions

In this study we reported results from the first proteomic investigation of the liver of heavy pigs using a label-free proteomic approach. Differences between the investigated breeds (Italian Duroc and Italian Large White) were derived by 25 proteins that resulted up-regulated in one or in the other groups of pigs. This work demonstrated that breed differences (underlying general genetic differences) might be highlighted at the liver proteome level that could be useful to describe breed-specific metabolic characteristics. We also indirectly provide evidences that quantitative proteomic approaches are useful to describe internal phenotypes that could be important to link and dissect external (production) and complex traits providing potential useful biomarkers in pig breeding programmes.

## Acknowledgments

We thank ANAS for the cooperation in collecting samples and personnel of the University of Bologna who helped in this activity. This study was supported by University of Bologna RFO 2017 funds, Italian MiPAAF (INNOVAGEN project).

The authors acknowledge the “Fondazione Cassa di Risparmio di Modena” for funding the HPLC-ESI-QTOF system at the Centro Interdipartimentale Grandi Strumenti (CIGS) of the University of Modena and Reggio Emilia.

## Supplementary Material

**S1 Table. Full list of the 200 proteins analysed in this study.**

**S2 Table. List of the 329 proteins used to group the liver proteins according to their different biological processes.**

**S3 Table. Links between differentially expressed proteins identified in the comparison between breeds and QTLs mapped in the corresponding gene regions (± 50 kbp).**

